# NeMO: a flexible R package for nested multi-species occupancy modeling and eDNA study optimization

**DOI:** 10.1101/2025.05.23.655794

**Authors:** Bastien Macé, Stéphanie Manel, Alice Valentini, Mathieu Rocle, Nicolas Roset, Erwan Delrieu-Trottin

**Affiliations:** CEFE, Univ Montpellier, CNRS, EPHE-PSL University, IRD, Montpellier, France; SPYGEN, Le-Bourget-du-Lac, France; Institut Universitaire de France, Paris, France; Compagnie Nationale du Rhône, Lyon, France; Office Français de la Biodiversité, Lyon, France

**Keywords:** Bayesian modeling, detection, elusive species, environmental DNA, false negatives, metabarcoding

## Abstract

Biodiversity monitoring using environmental DNA (eDNA) metabarcoding has expanded rapidly, providing a non-invasive tool widely adopted by ecologists and stakeholders. However, eDNA surveys are prone to imperfect detection, and non-detections are often misinterpreted as true absences – a critical issue when monitoring rare or elusive species. Despite its implications for biodiversity assessments, detection uncertainty is rarely quantified in eDNA-based studies. Occupancy modeling offers a powerful solution to this limitation but remains underused, partly due to a lack of accessible and flexible tools. We developed `NeMO` (Nested eDNA Metabarcoding Occupancy), a user-friendly R package for fitting multi-species occupancy models in a Bayesian framework. `NeMO` explicitly accounts for the nested structure of eDNA metabarcoding workflows – typically involving multiple replication steps such as field samples and PCR replicates – while accommodating presence/absence or read-count data. The framework estimates species occupancy, eDNA collection probability, amplification probability, and expected read counts, and allows users to assess the influence of environmental or methodological covariates on each process. Crucially, `NeMO` helps rigorously assess detectability and optimize resource allocation in eDNA surveys. It estimates the minimum number of eDNA samples, PCR replicates, and sequencing depth required to reliably detect species when present, thereby guiding study design. We illustrate its utility using a fish biodiversity dataset from the Rhône River (France). NeMO integrates key modelling features into a single streamlined framework, providing researchers and practitioners with an accessible and effective tool to assess detectability and optimize resource allocation in eDNA metabarcoding surveys. Our results highlight the importance of quantifying detection uncertainty, which has major implications for conservation monitoring and for designing cost-effective and reliable eDNA strategies.

**GRAPHICAL ABSTRACT:** `NeMO` provides a flexible framework for modeling multi-species site occupancy from eDNA data, estimating species presence and detectability despite imperfect detection. It supports diverse experimental designs, enables model comparison, and guides study design optimization by calculating minimal sampling effort to detect species, helping ecologists balance detection accuracy and resource allocation.

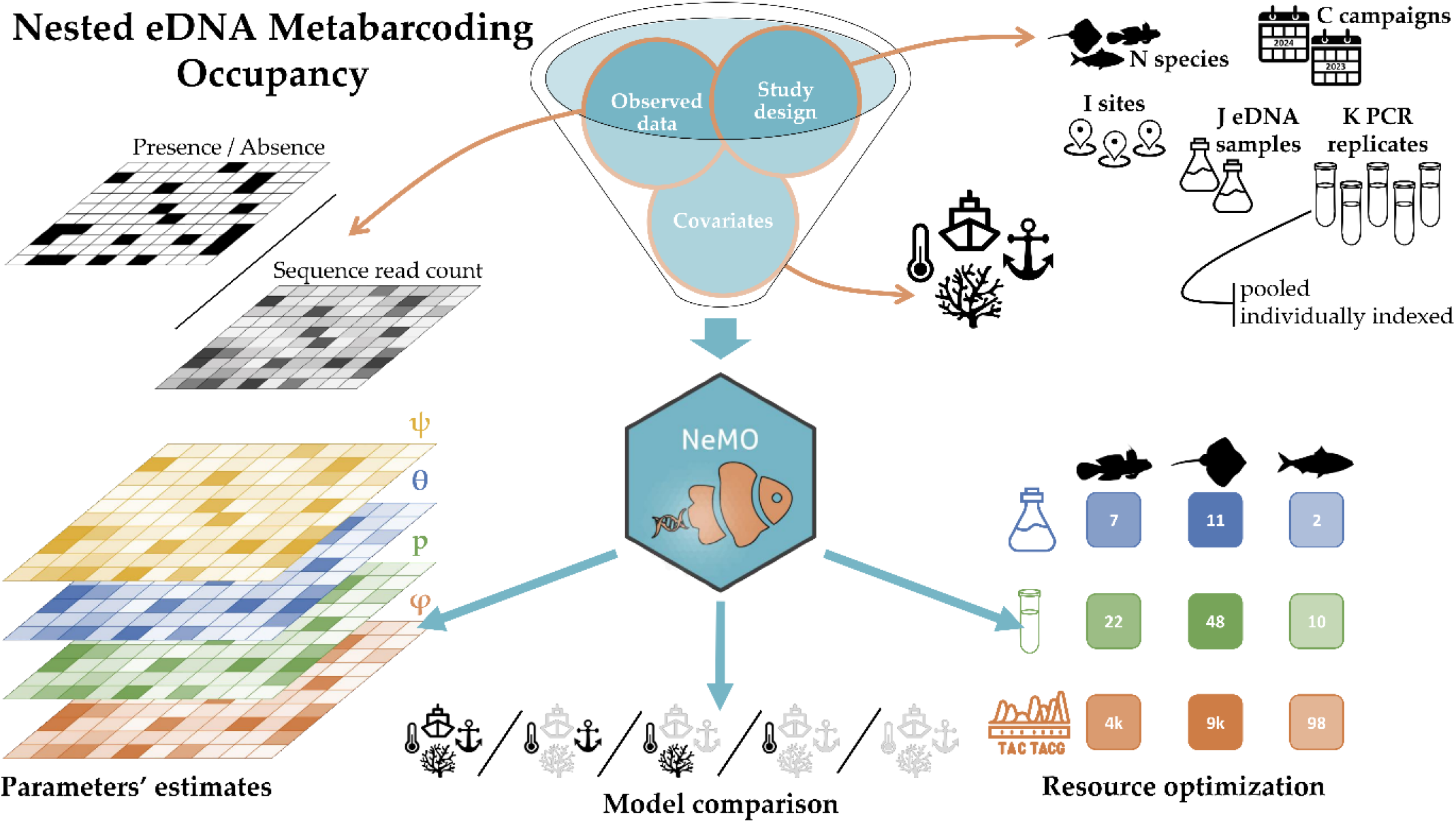

## 1. INTRODUCTION

Environmental DNA (eDNA) metabarcoding has emerged as a powerful tool for biodiversity monitoring (Blackman *et al*., 2024). Unlike traditional survey methods, which often rely on direct observations or specimen collection, eDNA metabarcoding enables the detection of a wide range of species from environmental samples such as soil, water, or air (Taberlet *et al*., 2012). This approach facilitates biodiversity assessments at the community level, providing a comprehensive view of ecosystem composition (Macé *et al*., 2024; Stat *et al*., 2017), including elusive species that conventional surveys may overlook (Beng & Corlett, 2020; Bohmann *et al*., 2014).

Like any biodiversity assessment approach, eDNA metabarcoding can be prone to detection errors (Cristescu & Hebert, 2018; Darling & Mahon, 2011), with false positives (*i*.*e*., species detected despite being absent from the sampled environment) and false negatives (*i*.*e*., species not detected despite being present in the sampled environment). False positives can result from contaminations, PCR or sequencing errors (Ficetola *et al*., 2016), or misidentifications due to incomplete or inaccurate reference databases (Polanco F. *et al*., 2025). To mitigate this, stringent laboratory protocols (Ficetola *et al*., 2016), including extraction blanks, PCR and sequencing controls, and bioinformatic filtering pipelines (Mathon *et al*., 2021), are routinely implemented. Various factors contribute to false negatives, such as low DNA concentration (Barnes *et al*., 2014), PCR primer biases during amplification (Alberdi *et al*., 2018), or insufficient sampling effort (Ficetola *et al*., 2015). Yet, unlike false positives, false negatives are often overlooked in eDNA studies. Absence data is frequently interpreted as true absences, without considering the possibility of undetected presence – an oversight that carries significant ecological implications (Biggs *et al*., 2015; Pinfield *et al*., 2019). Indeed, undetected species may lead to underestimated invasion dynamics, inadequate conservation efforts, or distorted community composition assessments.

Occupancy modeling offers a statistical framework to account for imperfect detection by estimating both occupancy and detection probabilities (MacKenzie *et al*., 2006). Originally developed for traditional field surveys (MacKenzie *et al*., 2002; Royle & Dorazio, 2008) relying on direct or indirect censuses, occupancy models have since been adapted for eDNA-based studies. These typically follow a hierarchical structure, incorporating multiple nested levels of replication (Takahashi *et al*., 2023). Specifically, the spatial units under investigation (*i*.*e*., sites) are generally sampled through different field replicates (*i*.*e*., eDNA samples), with each of these analyzed using multiple PCR replicates. Recognizing this structure, Schmidt *et al*. (2013) introduced the first nested occupancy model for single-species eDNA studies, estimating occupancy probability alongside detection probabilities at both the eDNA sample and PCR replicate levels. Building on this foundation, Dorazio & Erickson (2018) refined the approach in their `eDNAoccupancy` R package. Although this package improves accessibility and flexibility, the routine application of occupancy models remains limited in eDNA research, especially in multi-species metabarcoding studies, where non-detections are still frequently interpreted as true absences.

To extend the capabilities of existing models, occupancy frameworks tailored to eDNA metabarcoding studies have recently been introduced (**Table 1**), adapting the multi-species model from Dorazio & Royle (2005). While most of these models were made publicly available as text files, they were not initially designed as user-friendly software, which limited their accessibility to the broader scientific community. A major step forward was made with the development of dedicated R packages for multi-species occupancy modeling in eDNA studies. Fukaya *et al*. (2022) and Diana *et al*. (2024) introduced the `occumb` and `eDNAPlus` packages, respectively, both of which also integrating an important methodological advancement, *i*.*e*., the use of sequence read counts instead of solely presence/absence data. By incorporating quantitative sequencing information, these models refine detection probability estimates while keeping some limitations: `occumb` does not account for PCR replication while both are not allowing model comparisons. Building upon these advances, we introduce `NeMO` (Nested eDNA Metabarcoding Occupancy), a versatile R package designed to fully capture the complexity of eDNA metabarcoding datasets. `NeMO` accommodates a broad range of data structures, whether sequence read counts are available or only presence/absence data is provided. It also offers flexibility in handling different PCR replication schemes, whether replicates are individually indexed or pooled before sequencing. By integrating these features, `NeMO` enables users to choose the most suitable model based on the structure and requirements of their dataset. Like the previous occupancy modeling packages, `NeMO` allows the inclusion of covariates at multiple levels, enabling users to assess how environmental and methodological factors influence occupancy and detection probabilities. Additionally, we implement a model comparison tool, a feature present in `eDNAoccupancy` but not yet incorporated into multi-species occupancy frameworks. One key application of occupancy models is optimizing resource allocation in eDNA studies by estimating the minimum number of eDNA samples and PCR replicates needed to achieve a high probability of detecting species when present, thus explicitly accounting for imperfect detection. `NeMO` facilitates these calculations using equations from McArdle (1990) and extends this functionality by incorporating the option to estimate the minimum sequencing depth required for reliable species detection.

**Table 1.**
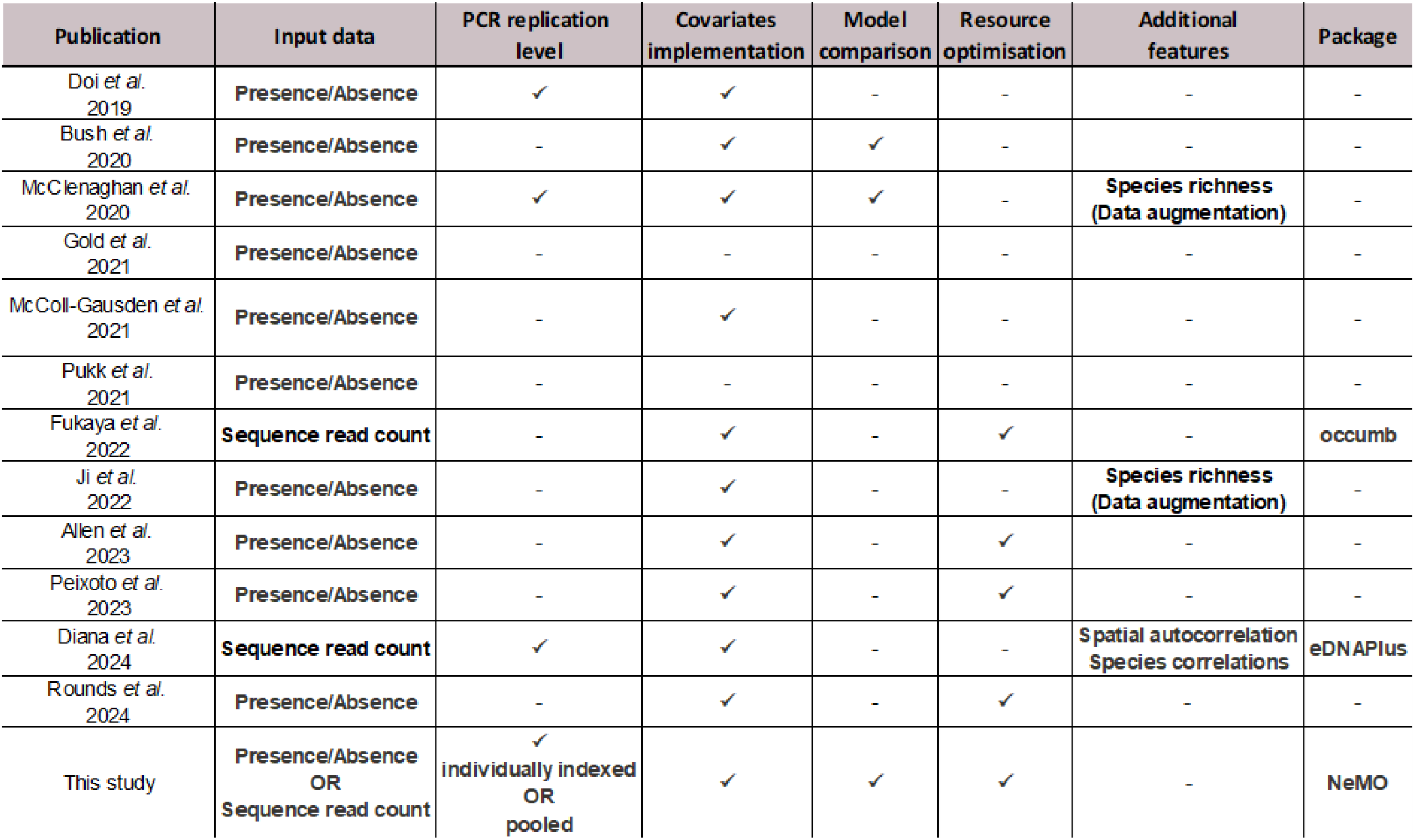
Scientific papers reporting code or software for multi-species occupancy modeling applied to eDNA metabarcoding, among the 75 papers identified through a literature search using the following keywords: [(“metabarcoding” OR (“eDNA” AND (“multispecies” OR “multi-species”))) AND “occupancy”] (Web of Science Core Collection on March 13, 2025). Full references are provided in **Appendix S3**.

To illustrate `NeMO` capabilities, we apply it to an empirical dataset from Pont *et al*. (2018), demonstrating its usefulness for ecological applications. Our goal is to provide an accessible, flexible, and powerful tool for scientists and stakeholders working with eDNA data.

## 2. MATERIAL & METHODS

### 2.1. Description of the nested eDNA metabarcoding occupancy model

eDNA-based surveys are typically conducted at multiple sites, with replication achieved by collecting several eDNA samples per site and running multiple PCRs for each sample. eDNA metabarcoding study designs therefore encompass three nested levels: site, eDNA sample, and PCR replicate (**Fig. 1a**). Additionally, high-throughput sequencing (HTS), which is commonly used to sequence eDNA barcodes (Taberlet *et al*., 2018), allows generation of sequence read counts for each produced amplicon (*i*.*e*. DNA fragment from a specific target region that has been amplified and sequenced), and ultimately for each species detected, provided that sequences are successfully assigned. Building on this framework, we adapted models proposed in Dorazio & Erickson (2018) and Fukaya *et al*. (2022) to fit multi-species occupancy and estimate parameters related to i) the probability of species eDNA occurrence at a site, ii) the conditional probability of collecting eDNA from a species in a sample from a site, given that its eDNA is present at the site, iii) the conditional probability of amplifying eDNA from a species in a PCR replicate of a sample, given that its eDNA is collected in the sample, and iv) the expected sequence read count generated for the species. To ensure consistency with existing literature, we adhere as closely as possible to the mathematical notations introduced by Schmidt *et al*. (2013) and Fukaya *et al*. (2022) for occupancy modeling in the context of the nested eDNA framework.

**Figure 1.**
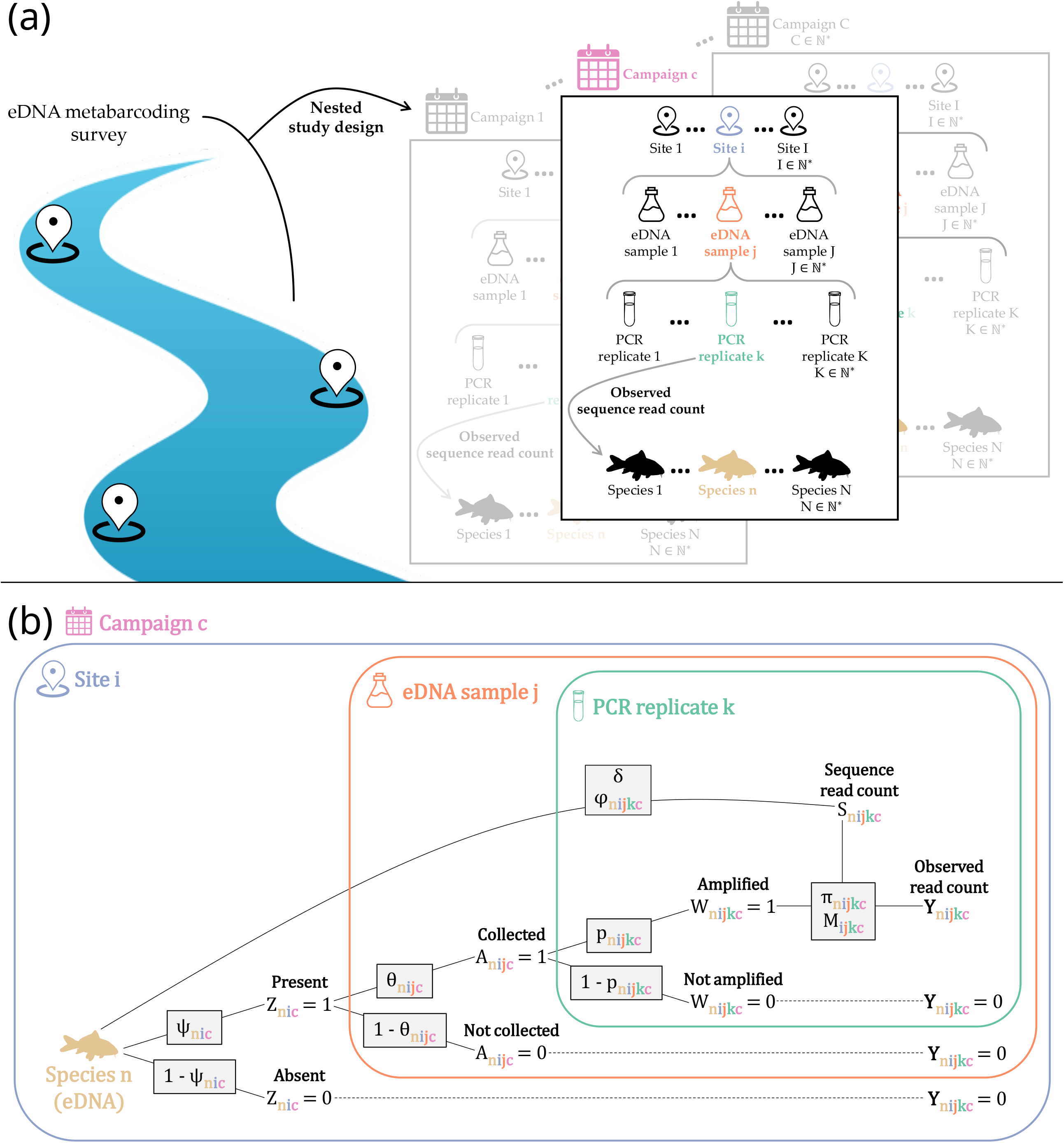
**(a)** Illustration of a nested study design for an eDNA metabarcoding survey in a river, monitoring *N* fish species over *C* campaigns, with replication across sites (*I*), samples (*j*), and PCR replicates (*K*). **(b)** Tree diagram illustrating the nested structure of the eDNA metabarcoding occupancy model. During a given campaign *c*, eDNA from species *n* may either be present at site *i* with occupancy probability *Ψ*_*nic*_ (latent variable *Z*_*nic*_ = 1), or absent with probability 1 − *Ψ*_*nic*_ (*Z*_*nic*_ = 0). If present, eDNA can be collected in an eDNA sample *j* with detection probability *θ*_*nijc*_ (latent variable *A*_*nijc*_ = 1), or remain uncollected with probability 1 − *θ*_*nijc*_ (*A*_*nijc*_ = 0). If collected, eDNA can be amplified in a given PCR replicate *k* with detection probability *p*_*nijkc*_ (latent variable *W*_*nijkc*_ = 1), or remain unamplified with probability 1 − *p*_*nijkc*_ (*W*_*nijkc*_ = 0). If amplified, eDNA is sequenced with frequency *Π*_*nijkc*_ relative to the total sequence read count *M*_*ijkc*_, generating the observed sequence read count *Y*_*nijkc*_. When eDNA is absent, not collected, or not amplified, the read count is *Y*_*nijkc*_ = 0. The observed read count depends on the sequence read count (latent variable *S*_*nijkc*_) which follows a negative binomial distribution with mean φ_*nijkc*_ and dispersion parameter *δ*.

We consider a community of *N* entities (*N* ∈ ℕ^*^), referred to as species for convenience, but which may also refer to taxa, operational taxonomic units (OTUs), amplicons, etc. Biodiversity monitoring of this community was repeated over time across *C* campaigns (*C* ∈ ℕ^*^), covering *I*_*c*_ sites (*I*_*c*_ ∈ ℕ^*^), where *J*_*i*_ samples (*J*_*ic*_ ∈ ℕ^*^) were collected per site and divided into *K*_*ijc*_ PCR replicates (*K*_*ijc*_ ∈ ℕ^*^). For each PCR replicate, HTS library preparation generated separate sequence reads. In this framework, the occupancy of species *n* ∈ {1, …, *N*} at site *i* ∈ {1, …, *I*_*c*_} during campaign *c* ∈ {1, …, *C*} is represented by the binary latent variable *Z*_*nic*_ which indicates whether eDNA from the species is present (*Z*_*nic*_ = 1) or absent (*Z*_*nic*_ = 0). This follows a Bernoulli distribution:

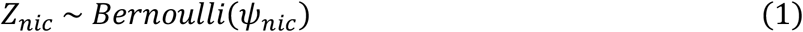

where *Ψ*_*nic*_ ∈ [0,1] is the probability that eDNA from species *n* being present at site *i* during campaign *c*. The detection of species *n* in sample *j* ∈ {1, …, *J*_*ic*_} from site *i* during campaign *c* is represented by the binary latent variable *A*_*nijc*_ which indicates whether eDNA from the species is collected (*A*_*nijc*_ = 1) or not (*A*_*nijc*_ = 0). This also follows a Bernoulli distribution, conditional on occupancy:

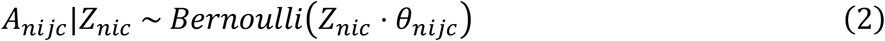

where *θ*_*nijc*_ ∈ [0,1] is the conditional probability of collecting eDNA from species *n* in sample *j* from site *i* during campaign *c*, given that species’ eDNA is present at site *i*. The detection of species *n* in PCR replicate *k* ∈ {1, …, *K*_*ijc*_} from sample *j* from site *i* during campaign *c* is represented by the binary latent variable *W*_*nijkc*_ which indicates whether eDNA from the species is amplified (*W*_*nijkc*_ = 1) or not (*W*_*nijkc*_ = 0). This also follows a Bernoulli distribution, conditional on eDNA collection:

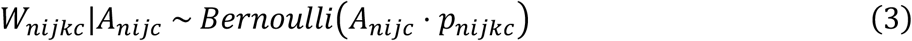

where *p*_*nijkc*_ ∈ [0,1] is the conditional probability of amplifying eDNA from species *n* in PCR replicate *k* from sample *j* from site *i* during campaign *c*, given that species’ eDNA was collected in sample *j*. The sequence read counts resulting from HTS are the observed variable *Y*_*nijkc*_. These counts are conditional on the per-PCR replicate total sequence read count *M*_*ijkc*_ ∈ ℕ hereafter referred to as ‘sequencing depth’ for simplicity. The sequencing depth is defined as:

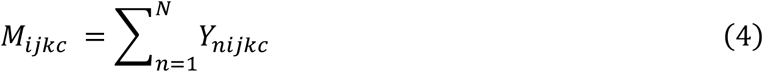

For *i, j, k, c* where *M*_*ijkc*_ > 0, the sequence read counts *Y*_*nijkc*_ follows a multinomial distribution, conditional on eDNA amplification:

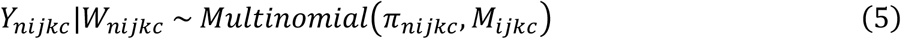

where *Π*_*nijkc*_ ∈ [0,1] is the conditional probability of assigning a sequencing read to species *n* in PCR replicate *k* from sample *j* from site *i* during campaign *c*, given that species’ eDNA was amplified in PCR replicate *k*. This parameter expresses the relative frequency of the sequence read count for species *n* under these conditions. It is defined as:

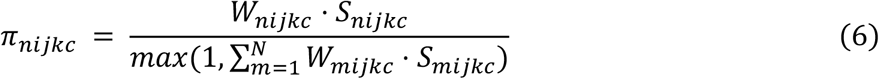

where the latent variable *S*_*nijkc*_ ∈ ℕ is the sequence read count for species *n* in PCR replicate *k* from sample *j* from site *i* during campaign *c*, and *W*_*nijkc*_ is the amplification detection indicator for the same replicate. The denominator ensures normalization of probabilities and accounts for cases where the product *W*_*mijkc*_ ⋅ *S*_*mijkc*_ is null (*e*.*g*., no amplification or zero read count for all species in the replicate). In such cases, the denominator defaults to 1. To accommodate the discrete and overdispersed nature of HTS data, we model the latent sequence read count *S*_*nijkc*_ using a negative binomial distribution:

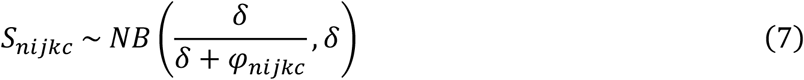

where φ_*nijkc*_ ∈ ℕ is the expected sequence read count for species *n* in PCR replicate *k* from sample *j* from site *i* during campaign *c*, and *δ* is the user-defined dispersion parameter controlling the variance of the distribution (defaults to 1). The framework of this occupancy model is summarized in **Fig. 1b**.

The species-level parameters *Ψ*_*nic*_, *θ*_*nijc*_, *p*_*nijkc*_, and *φ*_*nijkc*_ are modeled as random effects. Each parameter includes a species-specific intercept α and species-specific slope coefficients *β* associated with covariates that can be incorporated into the model. The linear predictors are defined as:

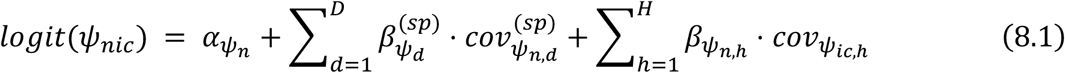

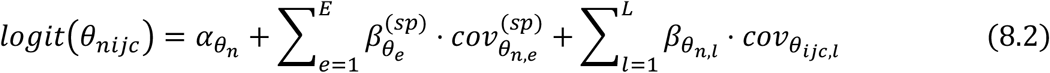

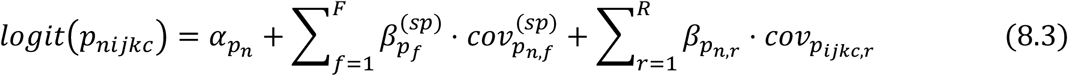

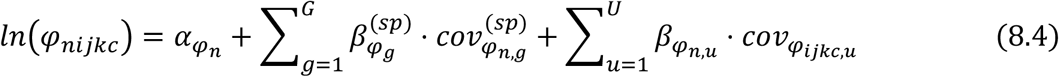

Here:

- 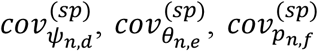, and 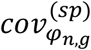 are Boolean or standardized species-level covariates associated with species *n*;
- 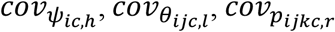, and 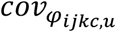 are Boolean or standardized spatial, methodological, or temporal covariate;
- *D, E, F, G* ∈ ℕare the numbers of species-level covariates;
- *H, L, R, U* ∈ ℕ are the numbers of spatial, methodological, or temporal covariates.

Species-level slopes associated with species covariates follow zero-centered normal priors:

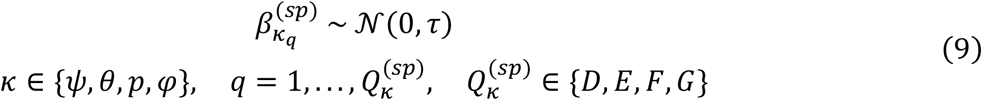

where *τ* > 0 is the user-defined hyperparameter controlling the precision (defaults to 1). Intercepts and slopes associated with spatial, methodological, or temporal covariates follow a multivariate normal community-level prior:

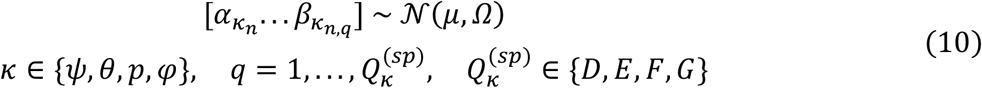

where μ is the community-mean vector and Ω the positive-definite community covariance matrix, capturing the variance of each parameter and the covariances between them. The community-mean vector μ follows a normal hyperprior:

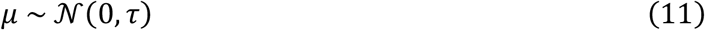

where *τ* > 0 is the user-defined hyperparameter controlling the precision (defaults to 1). The community covariance matrix Ω follows as Wishart hyperprior:

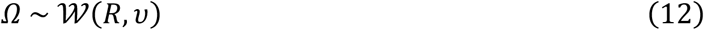

where *R* is a symmetric scale matrix with elements 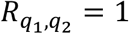 if 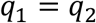 and 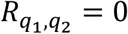 if *q*_1_ ≠ *q*_2_, determining the shape of the Wishart distribution, and *v* ∈ ℕ^*^ is the number of random effects.

Species are modeled as conditionally independent at the ecological levels (latent variables *Z, A, W, S*), but the associated parameters covary through a multivariate normal distribution (**Equation 10**), enabling borrowing of strength across the community. At the observation level (*Y*), the multinomial distribution creates negative correlations between species read counts due to the normalization and fixed sequencing depth. This compositional constraint reflects the technical reality of high-throughput sequencing, where species compete for a fixed pool of reads. Including species co-occurrence patterns in the modeling framework would probably provide a more detailed picture of species occupancy, but were not included here to avoid adding more ramifications to an already complex hierarchical model.

### 2.2. NeMO R package and associated functions

To analyze empirical data using our proposed model, we employed a Bayesian framework with Markov Chain Monte Carlo (MCMC) sampling, implemented with JAGS (Plummer, 2003). We developed a new R package, `NeMO` (Nested eDNA Metabarcoding Occupancy; available on GitHub: https://github.com/bastien-mace/NeMO), to facilitate this process. This package is designed to streamline the analysis of diverse eDNA metabarcoding experimental designs. Our proposed model comprehensively accounts for PCR replicates that are individually indexed for sequencing, a scenario that provides detailed information on presence/absence and sequence read count at the level of individual PCR replicates. However, pooling PCR replicates is a common strategy to reduce costs (Taberlet *et al*., 2018), but it limits the available information to summarized data. Additionally, while incorporating sequence read counts into occupancy modeling offers a more detailed approach, it also increases model complexity, which may not always be necessary depending on study objectives. To accommodate these variations, `NeMO` allows fitting an occupancy model tailored to the selected modeling protocol with the `Nemodel()` function:

1. ‘PCR_rep_seq_read’: This model represents the most complex scenario, incorporating eDNA samples, individually indexed PCR replicates, and sequence read counts.
2. ‘seq_read’: This model simplifies the analysis by omitting PCR replicates indexing (*k* level) while still considering sequence read counts. It modifies the model described above by replacing *W*_*nijkc*_ with *A*_*nijc*_ in **Equations 5-6** and excluding **Equation 3**, as well as the calculation of *p*_*nijkc*_ and associated random effects.
3. ‘PCR_rep’: This model focuses on individually indexed PCR replicates without incorporating sequence read counts. Here, *W*_*nijkc*_ is replaced with *Y*_*nijkc*_ in **Equation 3**, and **Equations 4-7** are excluded, as well as the calculation of *φ*_*nijkc*_ and associated random effects.

In addition to fitting occupancy models with the `Nemodel()` function, `NeMO` offers auxiliary tools to facilitate model implementation. It enables the integration of environmental or methodological covariates at any level of the nested model structure. The `covarray()` function standardizes these covariates into structured arrays compatible with the model, ensuring flexibility across study designs. The `min_resources()` function calculates the minimum number of samples, PCR replicates, and sequencing depth needed to achieve a specified probability of detecting species when present, *i*.*e*., to reliably detect species. The `WAIC()` function computes the Watanabe-Akaike Information Criterion (WAIC; Watanabe, 2010) for model comparison. These tools enhance `NeMO` by streamlining data preparation, guiding resource allocation, and aiding model selection. Detailed descriptions of these functions are available in **Appendix S1** and in the package documentation.

### 2.3. Application to empirical data

We modeled multi-species site-occupancy using a previously published dataset from an eDNA metabarcoding survey of fish conducted during a single campaign in 2016 along the Rhône River and its tributaries (Pont *et al*., 2018). 80 sites were selected to ensure a representative coverage of the entire river with a standardized study design (**Fig. 2a**), involving 2 eDNA samples per site and 12 PCR replicates per sample. Details of the sampling and laboratory methods are available in the original publication. The dataset includes sequence read count data for 100 fish species at the selected sites, including false positives – species absent from the Rhône River but detected with eDNA metabarcoding, likely resulting from human consumption or sewage contamination. Model fitting was performed using `NeMO` v1.0.0 with JAGS v4.3.2 (Plummer, 2003). To illustrate how covariates can be implemented, we included the distance to the sea as a site covariate at the occupancy level (*Ψ* parameter), as fish diversity gradients in the Rhône basin are well documented and particularly pronounced (Tockner *et al*., 2022). This dataset enabled us to run the `Nemodel()` function with the three different modeling protocols implemented, enabling comparison of model outputs. The raw sequence read count dataset was used for the ‘PCR_rep_seq_read’ protocol, while PCR replicates were pooled by sample for the ‘seq_read’ protocol, and sequence read counts were ignored and reduced to presence/absence data for the ‘PCR_rep’ protocol. Three MCMC chains were run in parallel, each with 150,000 iterations, a burn-in period of 100,000 and a thinning of 10. Eventually, we kept 5,000 iterations per chain (15,000 in total). Convergence was assessed using traceplots and the Gelman-Rubin statistic 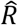 (Brooks & Gelman, 1998). The Genotoul-Bioinfo facility (Toulouse, France) was used to run and parallelize the MCMC chains across individual cores on standard nodes (128 threads, 2 TiB of RAM) equipped with dual AMD EPYC 7713 processors. The area under the curve (AUC) of the receiver operating characteristic (ROC) curve was calculated for each modeling protocol to compare model performance (Fawcett, 2006). This analysis requires prior knowledge of whether a species is present or absent at a given site, which was obtained using an exhaustive list of Rhône fish species supplemented by long-term electrofishing data (2006-2017) described in Pont *et al*. (2018). This dataset was restricted to the 10 years preceding the eDNA sampling (2007-2016). ROC analyses were conducted by river section to account for dams (**Fig. 2a**), using only sections with available electrofishing data. Species were classified as true positives, false negatives, true negatives, or false positives based on fish species list, eDNA and electrofishing detections (see **Appendix S2** for more details). The `pROC` R package v1.18.5 (Robin *et al*., 2011) was used to compute and plot ROC curves based on mean species occupancy probabilities from the latent *Z* array (**Equation 1**), providing a comprehensive assessment of model sensitivity and specificity.

**Figure 2.**
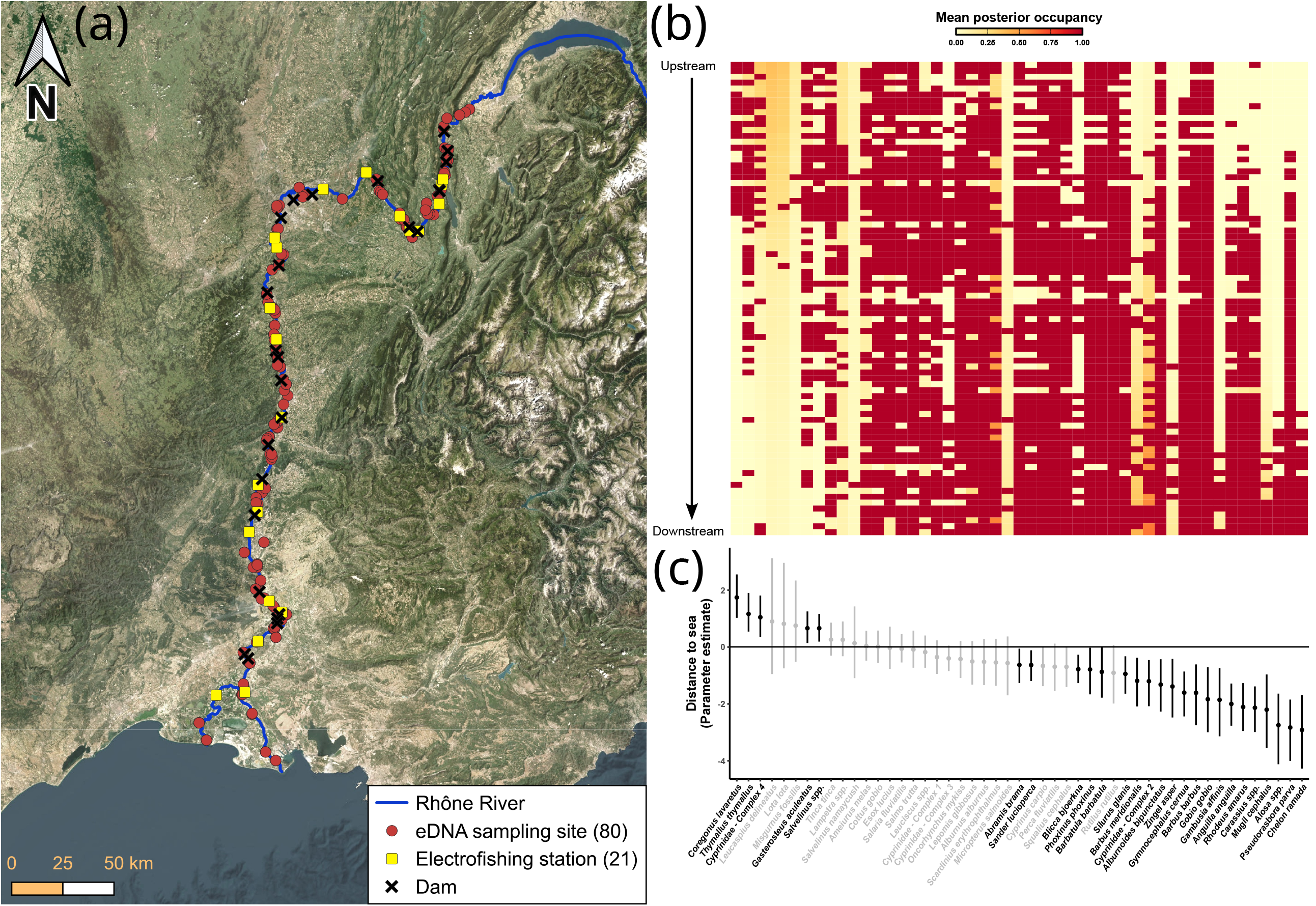
**(a)** Map of sampling locations along the Rhône River (France). The river thalweg is shown in blue. Red dots denote the 80 sites (including some located at the confluence of the Rhône and its tributaries) where two eDNA samples were collected per site. Yellow squares indicate stations where long-term electrofishing surveys were conducted, and black crosses mark the locations of dams. **(b)** Mean posterior occupancy (*Z* array) for species inhabiting the Rhône River at the 80 sampling sites. **(c)** Distance-to-sea occupancy parameter estimate (*β*_*Ψ*_) for each species inhabiting the Rhône River. Dots indicate the median estimates and their 95% HDI. Species with estimates whose HDI does not overlap zero are shown in black, others are shown in gray. Species identified at the genus level (*spp*.) or as part of complexes are detailed in **Table S2**.

Spearman’s rank correlation coefficients were calculated between the parameters’ link functions to examine whether the species eDNA occupancy probability (*Ψ*) is linked to its probability of collection (*θ*), amplification (*p*), and to its expected sequence read count (*φ*). For species that were recorded in the electrofishing dataset, correlations were also calculated between these parameters and the mean number of catches per species in the Rhône sections prospected over the ten-year survey period. This analysis aimed to investigate whether eDNA occupancy, collection, amplification and expected read count reflect quantitative abundance data. To determine the minimum resources required to reach a 95% detection probability (minimum number of samples *J*_*min*_, minimum number of PCR replicates *K*_*min*_, and minimum sequencing depth *M*_*min*_), we used the `min_resources()` function from the `NeMO` package presented here.

Analyses were conducted using R v4.3.2 (R Core Team, 2023). Unless otherwise stated, values are reported as medians with their 95% highest density intervals (HDIs) across the 15,000 MCMC samples ran with the ‘PCR_rep_seq_read’ protocol.

## 3. RESULTS

### 3.1. Protocol comparison & Model outputs

Model performance was similar across the three modeling protocols, with ROC analysis yielding comparable AUC values, ranging from 0.797 for ‘PCR_rep_seq_read’ to 0.824 for ‘seq_read’ (**Fig. S1**). Estimates remained consistent across all species regardless of the protocol used. However, execution time varied substantially (**Fig. S1**). ‘PCR_rep’ was the fastest, completing in just 2 hours with the high-performance computing facility, whereas ‘PCR_rep_seq_read’, the most computationally intensive design, required nearly 10 days to run on the same system. Given the similarity in model outputs across protocols, we focused further analyses on ‘PCR_rep_seq_read’ results, as this approach provides information at every level of the nested design.

Posterior occupancy for each species at all sites was obtained from the model output by extracting the latent *Z* array (**Fig. 2b**). Similarly, posterior latent matrices for collection (*A*), amplification (*W*), and read count (*S*) can be derived, though their visualization is challenging, as it would require three or more dimensions. Mean occupancy was generally very high when the species was detected with eDNA (close to 1) and very low otherwise (close to 0). However, upstream-downstream occupancy gradients were evident for some species, with certain taxa showing higher occupancy probability downstream, even in sites where they were not detected (*e*.*g*., *Barbus meridionalis, Cyprinidae – Complex 2*). These occupancy patterns align with the estimated effect of the distance-to-sea covariate (*β*_*Ψ*_, **Fig. 2c**). Based on this estimate, some species can be clearly identified as restricted to upstream sites (*β*_*Ψ*_ > 0: *e*.*g*., *Coregonus lavaretus, Thymallus thymallus*) or downstream sites (*β*_*Ψ*_ < 0: *e*.*g*., *Alosa spp*., *Chelon ramada*).

### 3.2. Correlations between parameters

Model outputs provided posterior probabilities for estimates related to eDNA occupancy (*Ψ*), collection (*θ*), amplification (*p*), and expected read count (*φ*). The probability densities varied noticeably among species (**Fig. S2**). However, species with high eDNA occupancy probability also tended to exhibit higher detectability (*θ, p*, and *φ*). This trend is further supported by positive Spearman’s rank correlation coefficients between all parameter pairs (**Fig. S3**). Correlations were particularly strong among detection-related parameters – eDNA collection (*θ*), amplification (*p*), and sequencing (*φ*) – ranging from 0.881 (HDI: 0.828 – 0.926) between *logit*(*θ*) and *log*(*φ*) to 0.961 (HDI: 0.931 – 0.983) between *logit*(*θ*) and *logit*(*p*). This indicates that species expected to yield high sequence read count also have eDNA more likely to be successfully amplified in PCR replicates and collected in samples.

Although occupancy and detection parameters were also positively correlated, these relationships were slightly weaker, with coefficients ranging from 0.573 (HDI: 0.482 – 0.655) between *logit*(*Ψ*) and *log*(*φ*), to 0.64 (HDI: 0.541 – 0.72) between *logit*(*Ψ*) and *logit*(*p*). As a result, species with high occupancy probability (*i*.*e*., occurring in more sites) also exhibited higher detectability across samples, PCR replicates, and sequencing reads, and species with low occupancy probability also exhibited lower detectability. The majority of species, like *Rutilus rutilus* (high *Ψ, θ, p*, and *φ*) and *Misgurnus fossilis* (low *Ψ, θ, p*, and *φ*) closely followed the general pattern, while *Alosa spp*. and *Scardinius erythrophthalmus* diverged (**Fig. S3**), illustrating the distinction between rarity (low occupancy) and elusiveness (low detectability). *Alosa spp*. was rare throughout the river (low *Ψ*) but highly detectable (high *θ, p*, and *φ*), reflecting its restriction to downstream sites (**Fig. 2b-c**) where its eDNA was consistently detected. In contrast, *Scardinius erythrophthalmus* was common (high *Ψ*) but elusive (low *θ, p*, and *φ*). **Figure 3** further illustrates these differences, showing consistently high or low parameter values for *Rutilus rutilus* and *Misgurnus fossilis*, while *Alosa spp*. and *Scardinius erythrophthalmus* exhibit the rare-conspicuous and common-elusive patterns, respectively.

**Figure 3.**
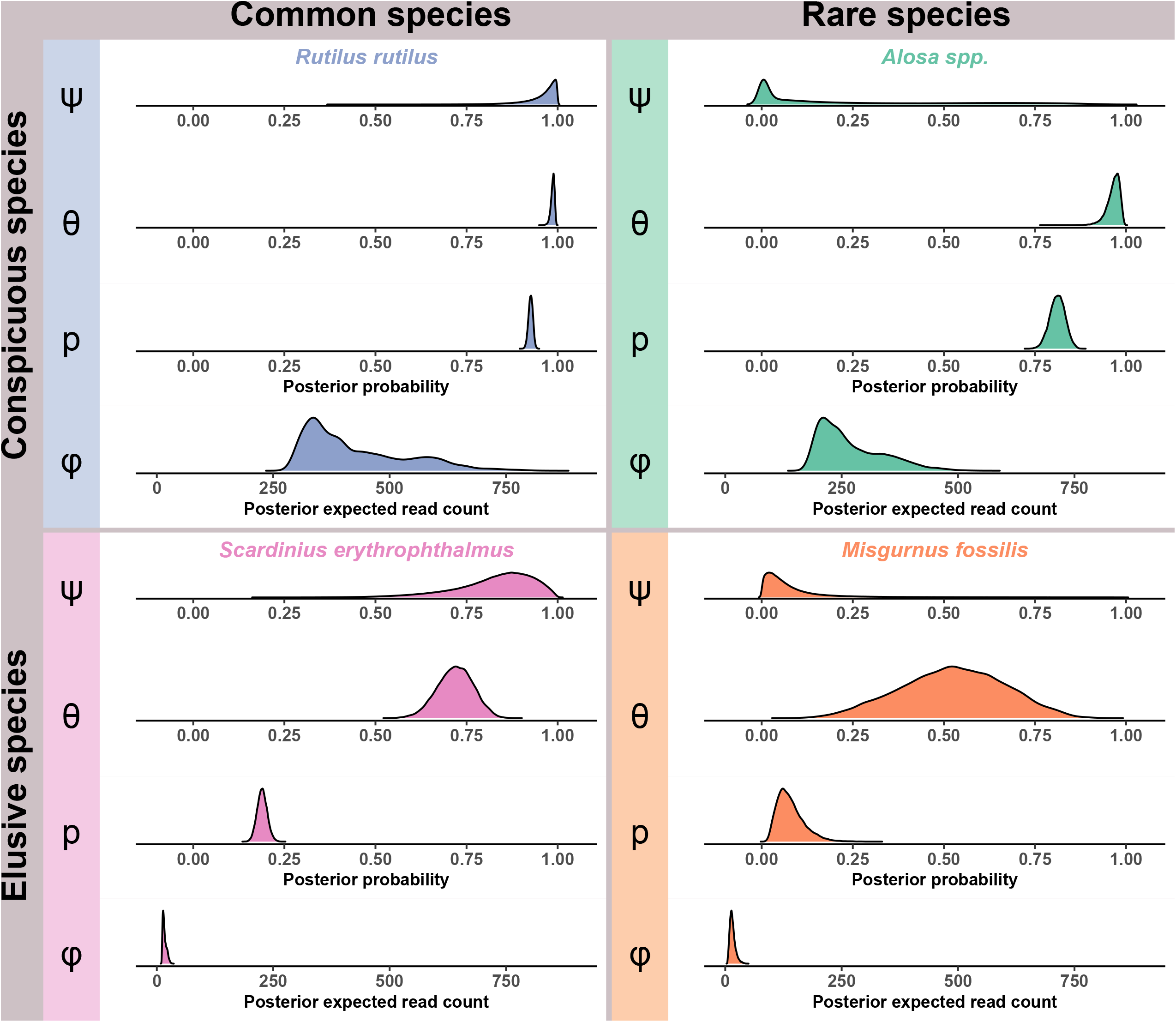
Ridge line plots depicting the densities of posterior probabilities for eDNA occupancy (*Ψ*), collection (*θ*), amplification (*p*), and expected read count (*φ*) for four species inhabiting the Rhône River. The highlighted species are *Rutilus rutilus* (blue), *Alosa spp*. (green), *Scardinius erythrophthalmus* (pink) and *Misgurnus fossilis* (orange). These species are categorized based on (1) their rarity – common species (*Rutilus rutilus* and *Scardinius erythrophthalmus*) have high occupancy probability; rare species (*Alosa spp*. and *Misgurnus fossilis*) have low occupancy probability – and (2) their detectability – conspicuous species (*Rutilus rutilus* and *Alosa spp*.) exhibit high eDNA collection and amplification probabilities, and have high expected read counts; elusive species (*Scardinius erythrophthalmus* and *Misgurnus fossilis*) are characterized by lower eDNA collection and amplification probabilities, as well as lower expected read count.

Spearman’s correlations were also positive when comparing the mean annual number of catches per species recorded during the ten-year electrofishing survey with parameters *Ψ, θ, p*, and *φ* (**Table S1**). They ranged from 0.4 (HDI: 0.377 – 0.421) for the expected read count (*φ*) to 0.516 (HDI: 0.456 – 0.571) for occupancy probability (*Ψ*). These results indicate a weak/moderate positive relationship between occupancy probability, detectability, and electrofishing-based abundance estimates.

### 3.3. Estimation of the minimum technical requirements (eDNA samples/PCR replicates/Sequencing depth) to reliably detect species

The minimum number of eDNA samples, PCR replicates, and minimum sequencing depth required to achieve a high probability of detecting species when present (95% detection probability) varied greatly among species (**Fig. 4**). Regarding eDNA sampling and PCR replication, our model shows that the effort reported in Pont *et al*. (2018) – *i*.*e*., 2 eDNA samples and 12 PCR replicates – was sufficient to reliably detect most species (63%). In contrast, 18 species would have required additional sampling and PCR technical effort to reach the same detection probability (potential false negatives). Among these, five species required just one additional sample, while they would all have required more PCR replicates. For some extreme cases, such as *Leucaspius delineatus* and *Lota lota*, the additional PCR replicates required would have been prohibitively high. The minimum sequencing required to reliably detect species was calculated for each sample, as individually indexed PCR products were pooled by sample in equal volumes during library preparation, targeting a theoretical sequencing depth of 500,000 reads (Pont *et al*., 2018). This estimated minimum sequencing depth was then compared to the observed sequencing depth, defined as the total number of reads per sample. Overall, sequencing depth was sufficient, as no samples fell below the required threshold for any species, except for *Leucaspius delineatus*, for which one sample at each of two locations did not reach the minimum required depth (**Fig. S4**). However, the second sample at each site still provided sufficient coverage to prevent detection failure.

**Figure 4.**
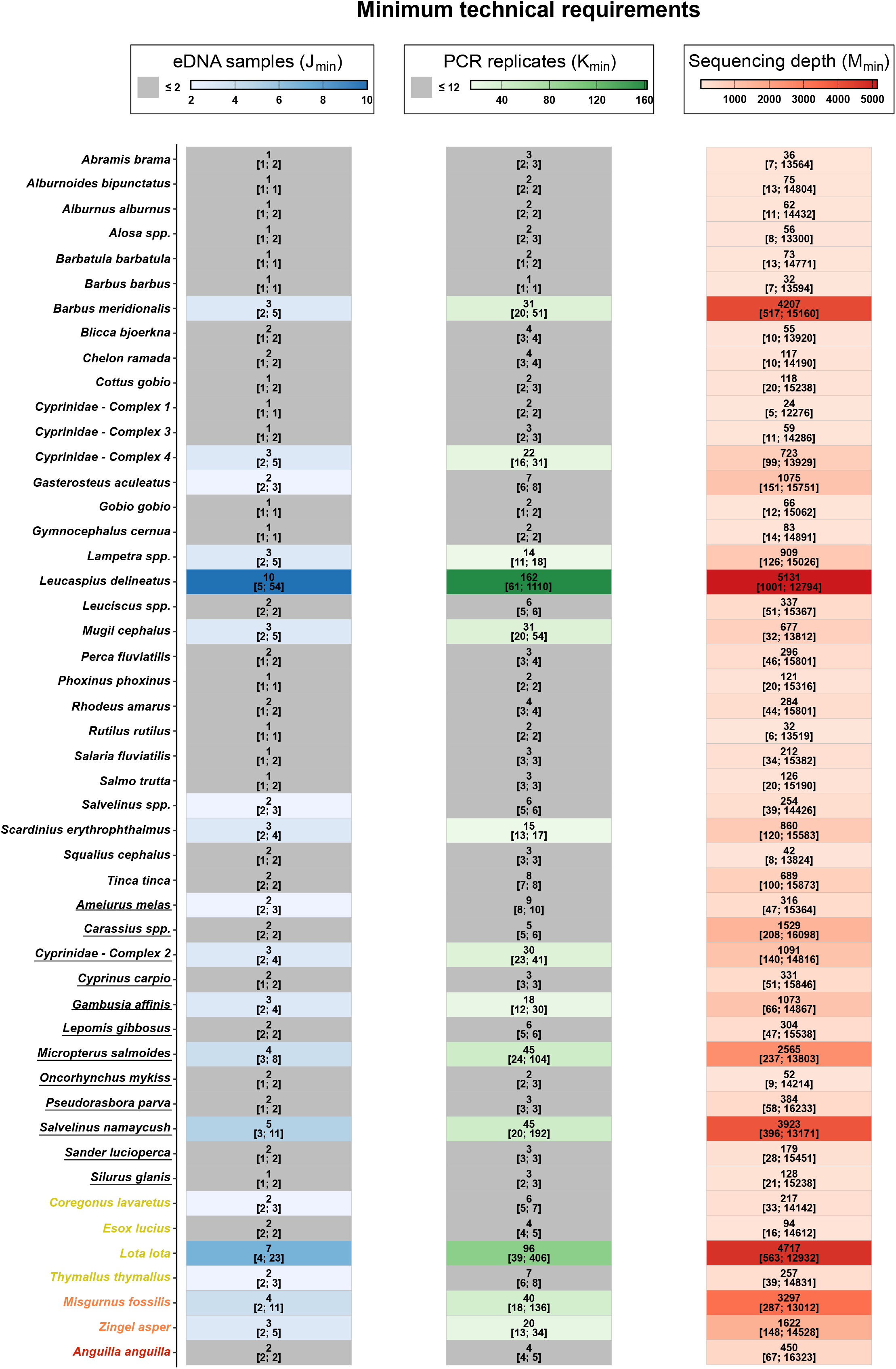
Tile plot illustrating the minimum number of eDNA samples (*j*_*min*_; in blue), PCR replicates (*K*_*min*_; in green), and sequencing depth (*M*_*min*_; in red) required to reliably detect species with 95% confidence. Median values are shown, with their 95% HDI in brackets. For sequencing, values are calculated across all samples and sites, providing summary estimates. The sequencing depth used in the study was always sufficient to detect species at all sites (see **Fig. S4**). For samples and replicates, gray shading indicates cases where the resources used in the original study (2 eDNA samples and 12 PCR replicates) were sufficient across the entire HDI. Species names are underlined if they are non-indigenous in the Rhône River, and threatened species are color-coded according to their IUCN conservation status at the most local scale possible (yellow: Vulnerable; orange: Endangered; red: Critically Endangered). Species identified at the genus level (*spp*.) or as part of complexes are detailed in **Table S2**.

## 4. DISCUSSION

We present `NeMO`, a Bayesian occupancy modeling framework tailored for metabarcoding-based eDNA studies with nested designs. This versatile R package accommodates diverse study designs, handling both presence/absence and sequence read count data, as well as individually indexed or pooled PCR replicates. `NeMO` accounts for multiple sources of uncertainty across hierarchical levels (collection, amplification, sequencing), and allows users to incorporate covariates influencing occupancy and detection probabilities. Beyond modeling, it enables formal model comparison to assess covariate effects and help optimize study design by evaluating whether sampling effort, PCR replication, and sequencing depth are sufficient to reliably detect species. By refining species occupancy estimates and guiding resource allocation, `NeMO` enhances the reliability of eDNA-based biodiversity assessments, particularly for elusive species.

We applied `NeMO` to a previously published eDNA metabarcoding dataset investigating fish diversity along the Rhône River (Pont *et al*., 2018). Regardless of the modeling protocol used, the high AUC (~ 0.8) in ROC analysis indicated strong model performance in distinguishing species presence from absence. By accounting for imperfect detection, occupancy probabilities provided more informative insights than raw presence/absence or sequence read count data. Integrating distance to sea as a covariate further refined site-level estimates, successfully capturing well-documented biological gradients (Keith *et al*., 2020) reflecting species distributions along the river system. Upstream-associated species had positive estimate values, including *Thymallus thymallus* and *Coregonus lavaretus*, the latter being restricted to alpine lakes. In contrast, downstream-associated species showed negative values, particularly diadromous species such as *Alosa spp*. (anadromous), *Anguilla anguilla* (catadromous), and also estuarine species such as *Chelon ramada*, reflecting their reliance on transitional or marine-influenced habitats.

Positive correlations between occupancy or detection parameter estimates with electrofishing data align with previous studies comparing eDNA with traditional metrics (Muri *et al*., 2020; Pont *et al*., 2018, 2023). These findings suggest that occupancy and detection probabilities could be meaningful proxies for species abundance in this river system. However, the correlations were moderate and these relationships should be interpreted cautiously when surveys differ in temporal and spatial scales. As expected, strong positive correlations were observed among detection parameters (*θ, p*, and *φ*), probably due to high eDNA abundance increasing the likelihood of successful collection in a sample, amplification in PCR replicates, and of obtaining a high number of sequencing reads. Correlations between detection parameters and occupancy probability (*Ψ*) were weaker, likely influenced by species-specific biological factors affecting DNA release into the environment, such as activity patterns, reproductive status, or body size (Stewart, 2019). Additionally, methodological factors like primer biases could affect DNA amplification, and sequencing efficiency despite species presence in the site surveyed (Elbrecht & Leese, 2015). Consequently, some species had low detectability and may remain undetected at sites they actually occupy. In this study, elusiveness (low detectability) is indeed particularly noticeable at the PCR level as *p* frequently displays low values and *φ* gets close to zero for many species (**Fig. S2**). While rare species are often elusive, exceptions exist: some common species remain difficult to detect (*Scardinius erythrophthalmus*), while certain rare species are conspicuous (*Alosa spp*.). This highlights the distinction between rarity and elusiveness, two concepts that should not be confused (Cerqueira *et al*., 2013). It is worth noting that these concepts should always be contextualized, as a species may be globally rare but regionally common – like upstream or downstream species in the Rhône River – and elusiveness is method-dependent (*i*.*e*., a species may be elusive because difficult to detect with one method but readily detectable with another).

`NeMO` enables retrospective evaluation of eDNA surveys by estimating the minimum number of eDNA samples, PCR replicates, and sequencing depth required to achieve a high probability of detecting species conditional on presence. While sequencing depth was consistently sufficient for the species surveyed by Pont *et al*. (2018), about one-third lacked enough samples or PCR replicates for reliable detection. This highlights the inevitable trade-off between resource allocation and species detectability. Although eDNA proved more effective than traditional methods for detecting elusive species – such as *Zingel asper, Mugil cephalus* and *Misgurnus fossilis*, which are elusive with eDNA metabarcoding but even more with electrofishing as they were never detected during the 10 years of survey – detectability must still be assessed before asserting its superiority. Our results suggest that increased replication effort could have allowed for the detection of these species at more sites. However, for some low-detectability species like *Leucaspius delineatus*, an ‘emerging’ species in the Rhône basin,achieving a high probability of detection when the species is present would require an impractical number of PCR replicates per sample (> 1,000), highlighting the limitations of the selected eDNA study design for certain targets. To improve detection, adjustments to study design could include alternative primers or modified sampling protocols. For example, *Lota lota*, a deep-dwelling nocturnal species (Keith *et al*., 2020), showed low detectability, likely because sampling was conducted in lentic systems at the surface and during daytime. Therefore, it is essential to cautiously interpret non-detections of such species, as it may not reflect true absence. We recommend integrating occupancy models into eDNA studies to validate non-detections and guide replication effort. This approach provides statistical support for filtering low-detection amplicons, as demonstrated by Pont *et al*. (2018), who excluded *Leucaspius delineatus* and *Lota lota* due to single-PCR detections – a decision now supported by `NeMO`. We further show that other species, such as *Salvelinus namaycush* or *Micropterus salmoides*, could have been filtered out as well. When eDNA sampling designs are insufficient for reliable species detection, practitioners face the challenge of deciding whether to increase the number of eDNA samples, PCR replicates, or sequencing depth. Ideally, minimum technical requirements should be met for all components. However, when budgetary or logistical constraints necessitate prioritization, we recommend the following approach: (i) identify which component(s) are most limiting for reliable detection based on study-specific objectives (*e*.*g*., targeted species, community completeness…); (ii) prioritize improvement of the component with the largest shortfall; and (iii) consider practical trade-offs. In most cases, increasing PCR replication is more cost-effective than adding field samples, as field sampling involves the entire downstream processing chain. However, PCR replication is constrained by the amount of extracted DNA available, and additional extractions can only be performed if biological material remains. Regarding sequencing depth, it is not freely adjustable as it is determined by the number of samples multiplexed on a sequencing run. Therefore, increasing either eDNA samples or PCR replicates automatically reduces read depth per sample unless additional sequencing runs are performed, which substantially increases costs. These considerations highlight the importance of using pilot studies to optimize sampling design in a statistically-driven way before downstream analyses. Given the risks of misinterpreting species absences, we recommend comparing community patterns with and without uncertain absences to assess their impact on α- and β-diversity, refining conclusions as necessary. Such practices can improve study efficiency and reliability of biodiversity assessments.

Among the three modeling protocols implemented in `NeMO`, the ‘PCR_rep’ protocol proved to be the fastest. When sequencing-related covariates are not included, models based on sequence read counts – such as the ‘seq_read’ and ‘PCR_rep_seq_read’ protocols in `NeMO –` could be avoided as they substantially increase computational time without improving performance. However, estimating the minimum sequencing depth required for reliable detection remains essential. For comparability, all models were run using the same number of MCMC iterations, which was chosen to ensure convergence of the most complex protocol, ‘PCR_rep_seq_read’. In practice, this indicates that a smaller number of iterations would likely have been sufficient for the two other protocols, further reducing their computational time relative to ‘PCR_rep_seq_read’. These analyses were conducted on a large empirical dataset, which contributes to the overall computation time. To facilitate testing and exploration of the workflow, the `NeMO` package includes a smaller example dataset derived from this dataset, allowing users to run the models and reproduce the main steps of the analysis within a few minutes. A dedicated vignette is also provided to guide users through the complete workflow using this example dataset.

While our framework accounts for false negatives and detection biases, it does not model false positives, which remain a common issue in eDNA studies. These are often mitigated through stringent bioinformatic pipelines or taxonomic filtering to exclude sequencing artifacts or unexpected detections arising from contamination or diet, which could also lead to false negatives if these filters are too stringent. Occupancy models that account for false positives do exist (Guillera-Arroita *et al*., 2017; Lahoz-Monfort *et al*., 2016), but they generally require additional data to distinguish true from false detections, such as an unambiguous detection method or calibration procedures applied at each stage of the multi-scale sampling hierarchy. These requirements add complexity to already intricate hierarchical models and are often impractical in large-scale metabarcoding studies. Additionally, in lotic systems, species detection may be spatially displaced due to downstream DNA transport, though this effect can be quantified (Perry *et al*., 2024; Pont *et al*., 2018). Future developments could thus integrate hydrological transport effects for both freshwater (Gimenez, 2024) and marine systems. More generally,incorporating spatial autocorrelation (Chen & Ficetola, 2019) or species co-occurrence (Rota *et al*., 2016) could improve inference, but at the cost of greater model complexity and computational demand.

## DATA AVAILABILITY STATEMENT

No new data were generated for this study. We used a previously published dataset available on Dryad (DOI: 10.5061/dryad.t4n42rr). The R package developed as part of this research, `NeMO`, is available on GitHub (https://github.com/bastien-mace/NeMO). The package includes a vignette and an example dataset to guide users in its implementation.

## ACKNOWLEDGEMENTS

We are grateful to the genotoul bioinformatics platform Toulouse Occitanie (Bioinfo Genotoul,https://doi.org/10.15454/1.5572369328961167E12) for providing computing and storage resources. We are grateful to Alexandre Granier for his insightful guidance on the structuring of our R package.

## AUTHOR CONTRIBUTION

SM and EDT conceived the study. BM analyzed the data and drafted the first manuscript. MR and NR provided essential data for the study. All authors contributed to manuscript revisions and approved the final version.

