## Supporting information for "NeMO: a flexible R package for nested multi-species occupancy modeling and eDNA study optimization"

---

### Contents

**Appendix S1.** Description of `NeMO`'s functions 2

1. **`Nemodel()`** 2
2. **`covarray()`** 4
3. **`min_ressources()`** 5
4. **`WAIC()`** 7

**Appendix S2.** Supplementary Methods 9

**Appendix S3.** Supplementary Figures & Tables 10

- **Figure S3.1** 10
- **Figure S3.2** 11
- **Figure S3.3** 12
- **Figure S3.4** 13
- **Table S3.1** 14
- **Table S3.2** 14

**Appendix S4.** Reference list of the papers retained from the literature research query 15

**References** 17

### **Appendix S1.** Description of `NeMO`'s functions

#### 1. **`Nemodel()`**

`Nemodel()` is the core of the `NeMO` package, designed for Bayesian modeling of multi-species occupancy in eDNA metabarcoding studies. It provides flexibility to fit different modeling protocols, accommodating various study designs and data structures. The function uses the `R2jags` R package (Su & Yajima, 2024) for MCMC sampling.

```
Nemodel(protocol = c('PCR_rep', 'seq_read', 'PCR_rep_seq_read'),
        array,
        covariates = NULL,
        name = 'model',
        nb_iterations = 2000,
        nb_burnin = floor(nb_iterations / 2),
        nb_thinning = max(1, floor((nb_iterations - nb_burnin) / 1000)),
        nb_chains = 3,
        parallel = FALSE,
        loglik = FALSE,
        latent = NULL,
        posterior = NULL,
        tau = 1,
        delta = 1,
        ...)
```

The modeling protocol is defined by the `protocol` argument, which determines the format of data input array (`array`) and the underlying model framework:

- `PCR_rep`: Requires a 5-dimensional ($N \times I \times J \times K \times C$) presence/absence input `array`.
- `seq_read`: Requires a 4-dimensional ($N \times I \times J \times C$) sequence read count input `array`.
- `PCR_rep_seq_read`: Requires a 5-dimensional ($N \times I \times J \times K \times C$) sequence read count input `array`.

The flexibility of this input design allows the user to specify data structures tailored to various experimental designs. The optional `covariates` argument allows users to include additional explanatory variables in the model. Covariates must be preprocessed using the `covarray()` function (see below). This feature allows for more nuanced modeling by accounting for external predictors. The following arguments control the MCMC sampling process:

- `name`: Filename and path of the BUGS language model file that will be recorded
- `nb_iterations`: Total number of iterations.
- `nb_burnin`: Number of initial iterations discarded as burn-in.
- `nb_thinning`: Interval of thinning to reduce autocorrelation in MCMC samples.
- `nb_chains`: Number of independent MCMC chains to run.

Parallel computation can be enabled by setting `parallel = TRUE`, which accelerates sampling for large datasets. Additional functionalities allow computation and saving of log-likelihoods (`loglik = TRUE`) for subsequent model comparison using `WAIC()` (see below), to save latent matrices (*e.g.*, $Z$, $A$, $W$, $S$, $Y$) with the `latent` argument, and to store posterior distributions of key parameters (*e.g.*, $\psi$, $\theta$, $p$, $\pi$, $\varphi$) using `posterior`. Recording these outputs is optional, allowing downstream analyses without recalculating them from the automatically recorded posterior distributions of the random effects. The `tau` argument ($\tau$, default: 1) controls the precision of normal (hyper)priors (**Equations 9, 11**). The `delta` ($\delta$, default: 1) argument correspond to the dispersion parameters of the negative binomial distribution (**Equation 7**). It should be set to high values (*e.g.*, $\delta$ = 10-50) for datasets with moderate overdispersion, and to low values (*e.g.*, $\delta$ = 0.1-5) for highly overdispersed data. Upon execution, the `Nemodel()` function fits the selected model to the provided array and generates a comprehensive class object with five primary slots:

- `model`: The fitted model output in `rjags` format (Plummer 2023). Additionally, a BUGS language text file is stored at the location specified by the `name` argument.
- `protocol`: The modeling protocol chosen.
- `array`: The input data array.
- `covariates`: The covariate list implemented in the model.
- `names`: Include the names of species, sites, samples, replicates and campaigns. This slot also contains the names of the recorded estimates, divided as follows:
    - `estimates_sp`: Names associated with the species-level random effects.
    - `estimates`: Names associated with the intercepts and other random effects.

2. **covarray()**

The covarray() formats covariates for integration into the NeMO modeling framework, producing arrays compatible with Nemodel().

```
covarray(protocol = c('PCR_rep', 'seq_read', 'PCR_rep_seq_read'),
         array,
         cov_list = list(
            list(cov_data = NULL,
                 level = 'psi',
                 dimension = 'species')))
```

The modeling protocol and input data array are given with the protocol and array arguments respectively, as described earlier. Covariates are given with the cov_list argument. They are provided as a list of sublists, each representing a covariate, and must have a unique name to facilitate identification and prevent conflicts. Sublists must include the following components:

- cov_data: An array containing covariate values. These values should be either Boolean for categorical/semi-quantitative predictors or standardised values for quantitative predictors.
- level: The hierarchical level where the covariate applies (*e.g.*, psi, theta, p, phi). A slope term associated with the covariate will be calculated at the selected level during MCMC sampling.
- dimension: Associated dimensions of the covariate. Valid values include:
    - Single dimensions: species, site, sample, replicate, or campaign.
    - Combined dimensions: Combinations of the above, except species, which always stands alone. In combinations, terms must be separated by an underscore '_', in the following order: 1) site 2) sample 3) replicate 4) campaign (*i.e.*, site_campaign).

The cov_data array in each sublist must align with the specified dimension. For instance, if dimension = 'site_sample_campaign', the covariate array should be structured such that dimension 1 corresponds to sites, dimension 2 corresponds to samples, and dimension 3 corresponds to campaigns. The covarray() function generates a structured class object with eight slots:

- psi_cov: Covariates for spatial/methodological/temporal components of $\psi$.

- `psi_cov_sp`: Species covariates of $\psi$.
- `theta_cov`: Covariates for spatial/methodological/temporal components of $\theta$.
- `theta_cov_sp`: Species covariates of $\theta$.
- `p_cov`: Covariates for spatial/methodological/temporal components of $p$.
- `p_cov_sp`: Species covariates of $p$.
- `phi_cov`: Covariates for spatial/methodological/temporal components of $\varphi$.
- `phi_cov_sp`: Species covariates of $\varphi$.

These slots organise the covariates across the four hierarchical levels, enabling their direct implementation into the modeling framework (refer to **Equation 9** for mathematical details).

##### 3. `min_resources()`

The `min_resources()` function calculates the minimal resource requirements – samples ($J_{min}$), PCR replicates ($K_{min}$), and sequencing depth ($M_{min}$) – needed to achieve a specified probability of detecting species when present (95% for example). This estimation relies on established probabilistic models, enabling rigorous resource planning for ecological studies.

```
min_resources(model,
              resources = c('J'),
              conf = 0.95)
```

The calculations underpinning the function follow equations derived from McArdle (1990) and standard probabilistic models:

- Minimum number of samples ($J_{min}$): Using **Equation 2**, and assuming samples are comparable and independent, we calculate the probability that DNA from species $n$ is not collected by the sample $j$ at site $i$ during campaign $c$, given that species DNA is present at site $i$, such that this probability is less than 0.05, corresponding to a 95% confidence level:

$$P(A_{nijc} = 0 | Z_{nic} = 1) \leq 0.05 \quad \Leftrightarrow \quad (1 - \theta_{nijc})^{J_{min_{nijc}}} \leq 0.05$$
$$\Leftrightarrow \quad J_{min_{nijc}} \geq \frac{ln(0.05)}{ln(1 - \theta_{nijc})}$$

- Minimum number of PCR replicates ($K_{min}$): Using **Equation 3**, and assuming PCR replicates are comparable and independent, we calculate the probability that DNA from species $n$ is not amplified in the PCR replicate $k$ from the sample $j$ at site $i$ during campaign $c$, while species' DNA is collected in sample $j$:

$$P\left(W_{nijkc}=0\middle|A_{nijc}=1\right)\leq 0.05 \quad \Leftrightarrow \quad \left(1-p_{nijkc}\right)^{K_{min_{nijkc}}}\leq 0.05$$
$$\Leftrightarrow \quad K_{min_{nijkc}}\geq \frac{ln(0.05)}{ln\left(1-p_{nijkc}\right)}$$

- Minimum sequencing depth ($M_{min}$): Using **Equation 5**, and following the multinomial theorem, we calculate the probability that DNA from species $n$ is not sequenced for the PCR replicate $k$ from the sample $j$ at site $i$ during campaign $c$:

$$P\left(Y_{nijkc}=0\middle|W_{nijkc}=1\right)\leq 0.05$$
$$\Leftrightarrow \sum_{y_{11111}+\dots y_{nijkc-1}+y_{nijkc+1}+\dots+y_{NIJKC}} \frac{M_{min_{nijkc}}}{y_{11111}!\dots y_{nijkc-1}!\,y_{nijkc+1}!\dots y_{NIJKC}!}\prod_{\varepsilon\neq nijkc}\pi_{\varepsilon}\leq 0.05$$
$$\Leftrightarrow \left(\sum_{\varepsilon\neq nijkc}\pi_{\varepsilon}\right)^{M_{min_{nijkc}}}\leq 0.05$$
$$\Leftrightarrow \left(1-\pi_{nijkc}\right)^{M_{min_{nijkc}}}\leq 0.05$$
$$\Leftrightarrow M_{min_{nijkc}}\geq \frac{ln(0.05)}{ln\left(1-\pi_{nijkc}\right)}$$

These computations are automatically handled within the function for precise and efficient resource estimation. The `model` argument is the `Nemodel` fitted model object, used to extract parameter estimates for resource calculations. Minimum resources to calculate are specified with the `resources` argument. Acceptable values are `J`, `K`, and `M`, and multiple resource types can be calculated simultaneously by supplying a vector (*e.g.*, `resources = c('J', 'K', 'M')`). Note that the `protocol` recorded in the `model` object determines which resource types can be computed:

- `J` can be computed for all protocols.
- `K` requires `PCR_rep` or `PCR_rep_seq_read`.
- `M` requires `seq_read` or `PCR_rep_seq_read`.

The probability to detect species when present, *i.e.*, confidence level, is defined with `conf` (default: 0.95). The `min_resources()` function generates a structured class object containing three slots: $J_{min}$, $K_{min}$, and $M_{min}$. Each slot corresponds to a list with the median value and the lower and upper bounds of the 95% highest density interval (HDI).

4. **`WAIC()`**

The `WAIC()` function computes the Watanabe-Akaike information criterion (WAIC), sometimes referred to as Widely Applicable Information Criterion (Watanabe, 2010). This robust metric is used for model comparison and selection within Bayesian frameworks, offering a way to balance model fit and complexity.

```
WAIC(model)
```

This calculation is conditional on having set `loglik = TRUE` in the `Nemodel()` function that produced the fitted `model` argument, ensuring that the log-likelihoods are recorded during model fitting. The WAIC metric is calculated using the following equations, which consider both the mean log-likelihood and its variance to quantify predictive accuracy and penalizing overfitting:

1. Log Pointwise Predictive Density ($LPPD$):

$$LPPD = \sum_{N}\sum_{I}\sum_{J}\sum_{K}\sum_{C} ln\left(\frac{1}{Sim_{tot}}\sum_{sim=1}^{Sim_{tot}} e^{log\_lik_{nijkc,sim}}\right)$$

where $log\_lik_{nijkc,sim}$ is the log-likelihood value calculated from simulation $sim$ for species $n$, at site $i$, in sample $j$, in replicate $k$ (except for `seq_read` protocol), during campaign $c$, and $Sim_{tot}$ is the total number of simulations (MCMC samples) computed.

2. Effective number of parameters ($p_{WAIC}$):

$$p_{WAIC} = \sum_{N}\sum_{I}\sum_{J}\sum_{K}\sum_{C} \frac{1}{Sim_{tot}}\sum_{sim=1}^{Sim_{tot}} \left(log\_lik_{nijkc,sim} - \overline{log\_lik_{nijkc,sim}}\right)^2$$

where the variance of $log\_lik_{nijkc,sim}$ around its mean $\overline{log\_lik_{nijkc,sim}}$ reflects the model's complexity by accounting for the variability in log-likelihood across posterior MCMC samples.

3. WAIC score:

$$WAIC = -2(LPPD - p_{WAIC})$$

These computations are performed internally, yielding a scalar WAIC value that summarizes the overall performance of the model. The `WAIC()` function returns a structured class object containing:

- `waic`: The WAIC score, quantifying the trade-off between fit and complexity.
- `lppd`: The log-predictive density term, reflecting the model's fit.

- `p_waic`: The effective number of parameters, representing model complexity.

This detailed output enables users to interpret both the predictive accuracy and the complexity of the fitted model.

### **Appendix S2.** Supplementary Methods

#### Application to empirical data

The area under the curve (AUC) of the receiver operating characteristic (ROC) curve was calculated for each modeling protocol to compare model performance (Fawcett, 2006). ROC curves are generated by plotting sensitivity (true positive rate, TPR) against 1 – specificity (false positive rate, FPR) at various threshold values. These thresholds range from 0 to 1 and represent the decision boundary for classifying species as present or absent. By varying this threshold, the model's performance across the full spectrum of classification criteria can be evaluated. Sensitivity is calculated as $TPR = \frac{TP}{TP+FN}$, where TP is the number of true positives, and FN the number of false negatives. Specificity is calculated as $TNR = \frac{TN}{TN+FP}$, where TN is the number of true negatives, and FP is the number of false positives. This analysis relies on prior knowledge of species presence or absence at each site, which was obtained using an exhaustive list of Rhône fish species, supplemented by long-term electrofishing data (2006-2017) described in Pont *et al.* (2018). This dataset was restricted to the 10 years preceding the eDNA sampling (2007-2016). ROC analyses were conducted by river section to account for dams (**Fig. 2**), using only sections with available electrofishing data. In every section, eDNA detections were classified as follows:

- True positive (TP): Species detected and known to inhabit the Rhône River.
- False negative (FN): Species not detected but known to inhabit the Rhône River and reported at least once in electrofishing surveys.
- True negative (TN): Species not detected and known to be absent from the Rhône River.
- False positive (FP): Species detected and known to be absent from the Rhône River.

The pROC R package v1.18.5 (Robin *et al.*, 2011) was used to compute and plot ROC curves based on mean species occupancy probabilities from the $Z$ array, providing a comprehensive assessment of model sensitivity and specificity.

### **Appendix S3.** Supplementary Figures & Tables

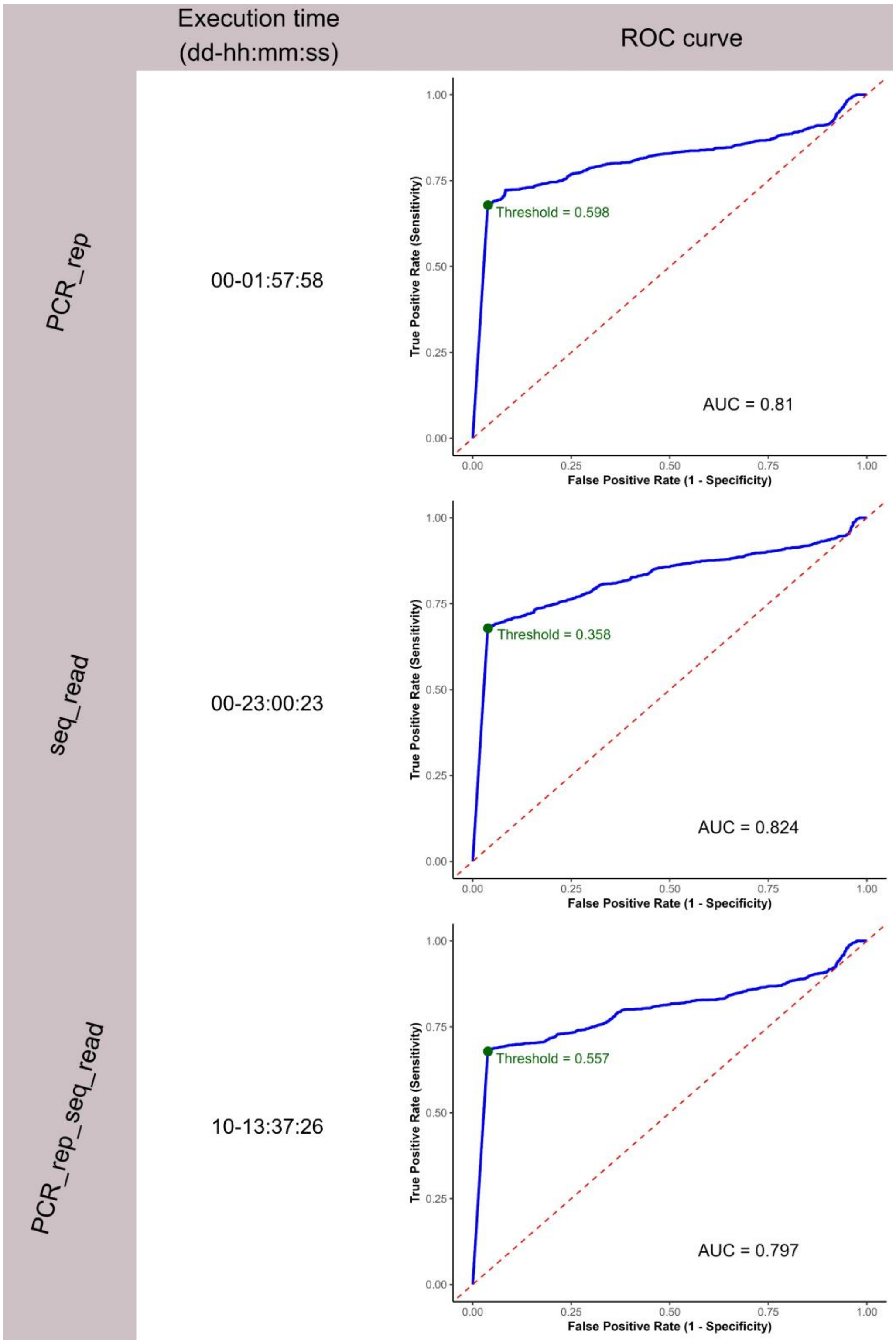


**Figure S3.1.** Comparison of the three protocols based on execution time (formatted as dd-hh:mm:ss), and ROC curves. AUC and optimal threshold values – determined using Youden's index (Youden, 1950) – are shown.

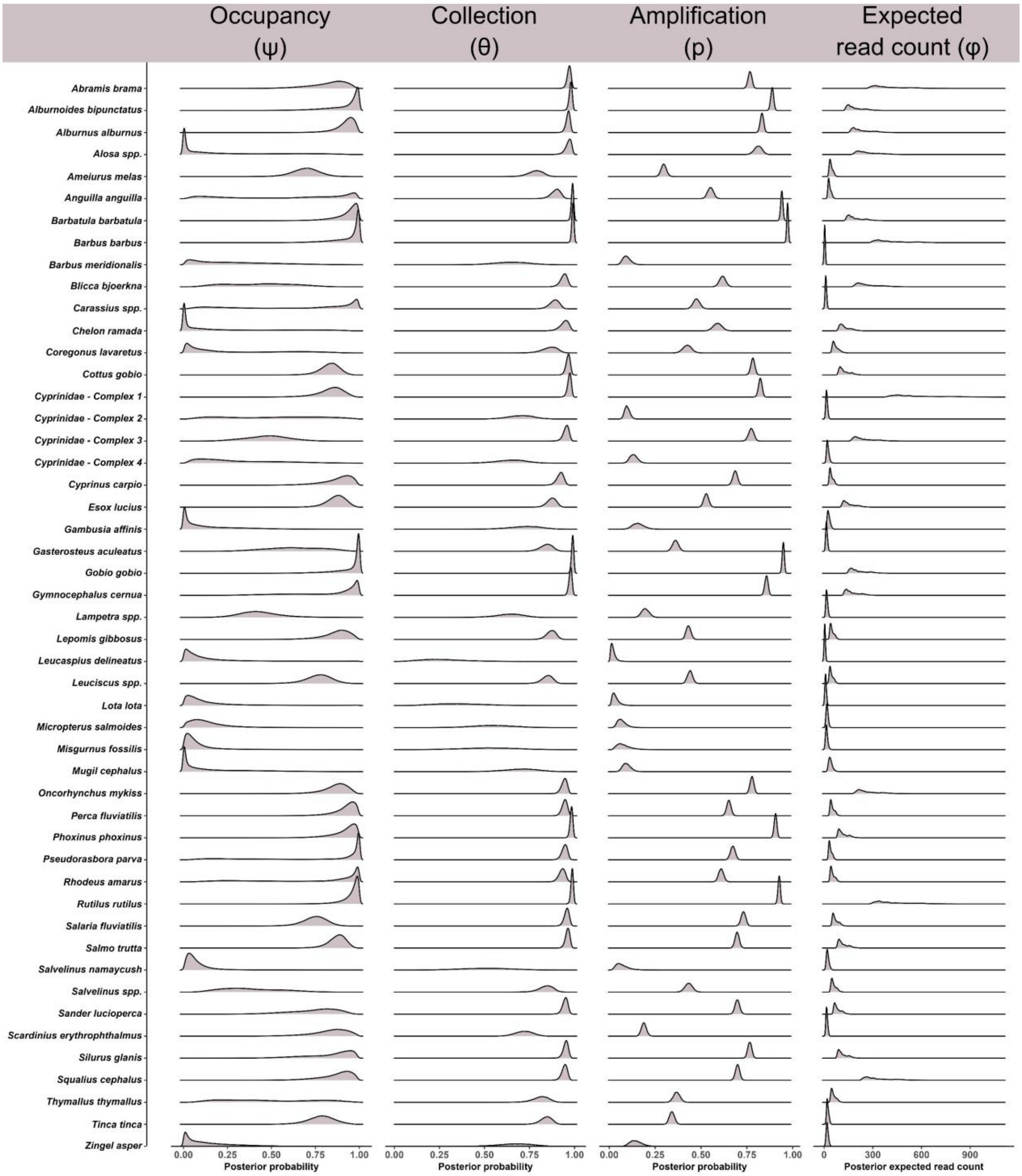


**Figure S3.2.** Ridge line plots depicting the densities of posterior probabilities for eDNA occupancy ($\psi$), collection ($\theta$), amplification ($p$), and expected sequence read count ($\varphi$) for species inhabiting the Rhône River. Species identified at the genus level (*spp.*) or as part of complexes are detailed in **Table S3.2**.

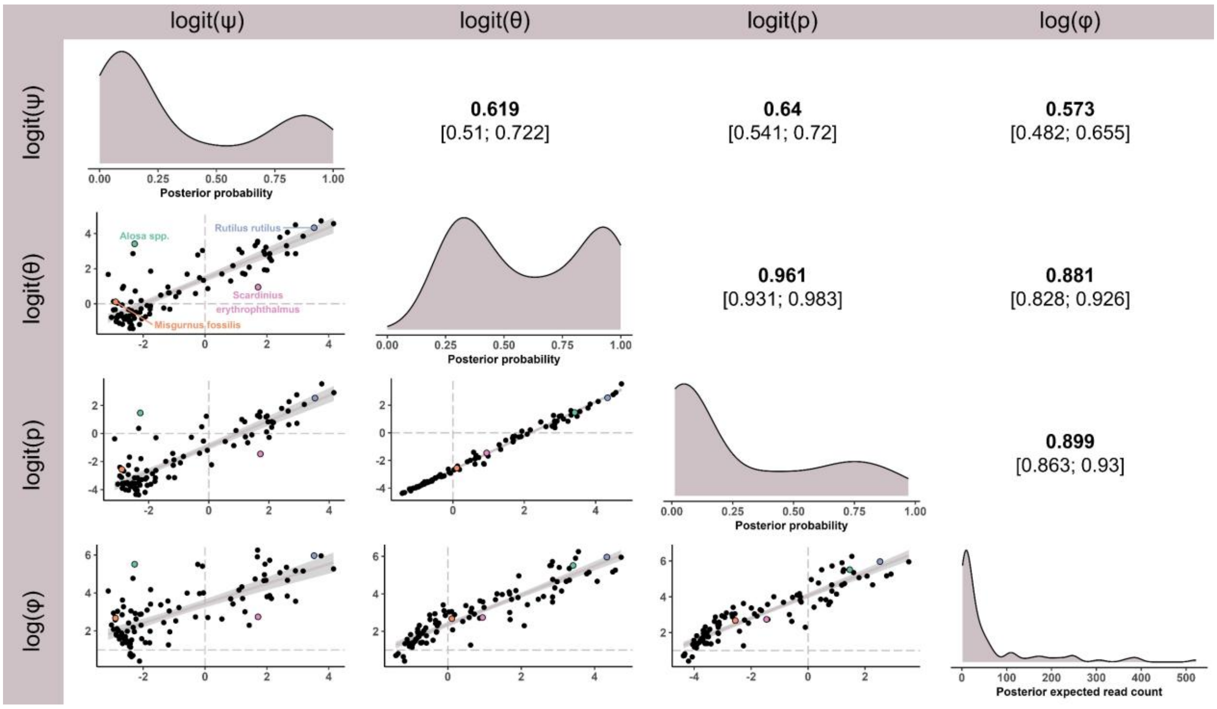


**Figure S3.3.** Upper triangular matrix: Median Spearman's rank correlation coefficients with their HDI in brackets, calculated between the link functions of parameters $\psi$ (occupancy), $\theta$ (collection), $p$ (amplification), and $\varphi$ (expected sequence read count) across all species. Diagonal: Posterior densities of parameter medians across all species. Lower triangular matrix: Scatter plots showing pairwise relationships between medians of parameter link functions across all species. Points are coloured for four species: *Rutilus rutilus* (blue), *Alosa spp.* (green), *Scardinius erythrophthalmus* (pink), and *Misgurnus fossilis* (orange).

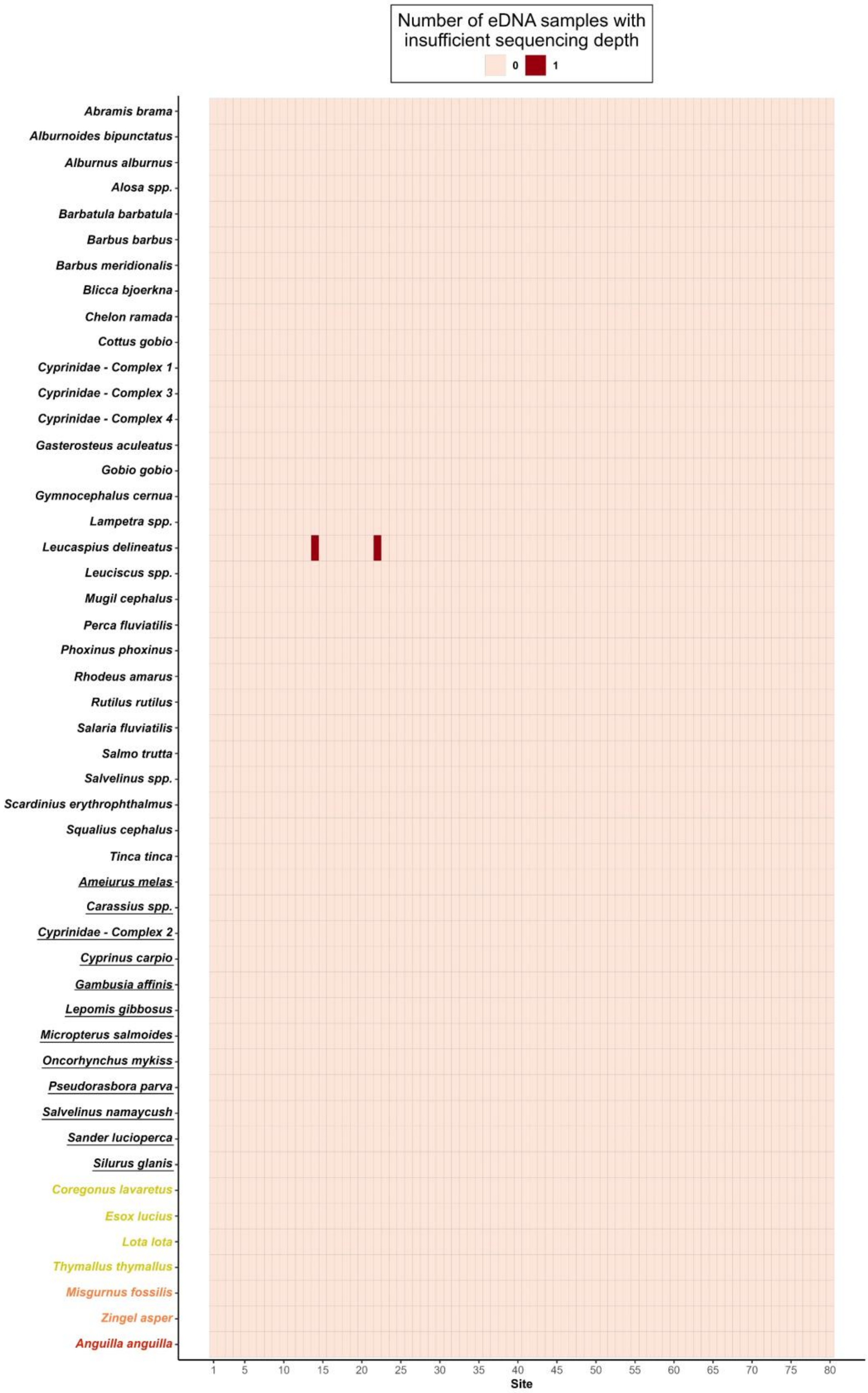


**Figure S4.** Tile plot showing the median number of samples per site with insufficient sequencing depth to detect species with 95% confidence when they are present. Species names are underlined if non-indigenous in the Rhône River, and threatened species are colour-coded according to their IUCN conservation status at the most local scale possible (yellow: Vulnerable; orange: Endangered; red: Critically Endangered). Species identified at the genus level (*spp.*) or as part of complexes are detailed in **Table S2**.

**Table S3.1.** Median Spearman's rank correlation coefficients with their HDI in brackets, calculated between the parameters $\psi$, $\theta$, $p$, $\varphi$ and the mean catch per unit effort (CPUE) for every species over the ten-year electrofishing survey.

| | **Occupancy (ψ)** | **Collection (θ)** | **Amplification (p)** | **Expected read count (φ)** |
|---|---|---|---|---|
| **Mean electrofishing CPUE** | **0.516** [0.456; 0.571] | **0.449** [0.382; 0.52] | **0.451** [0.438; 0.466] | **0.4** [0.377; 0.421] |

**Table S3.2.** List of species assigned to each genus or species complex in the dataset, where the DNA barcode sequence does not allow species-level discrimination. Species names are underlined if non-indigenous in the Rhône River, and threatened species are colour-coded according to their IUCN conservation status at the most local scale possible (yellow: Vulnerable; orange: Endangered; red: Critically Endangered).

| Genus / Complex | Species |
|---|---|
| ***Alosa spp.*** | *Alosa alosa*<br>*Alosa agone*<br>*Alosa fallax* |
| ***Carassius spp.*** | *Carassius auratus*<br>*Carassius carassius*<br>*Carassius gibelio* |
| ***Cyprinidae - Complex 1*** | *Chondrostoma nasus*<br>*Parachondrostoma toxostoma*<br>*Telestes souffia* |
| ***Cyprinidae - Complex 2*** | *Ctenopharyngodon idella*<br>*Hypophthalmichthys molitrix* |
| ***Cyprinidae - Complex 3*** | *Abramis brama*<br>*Blicca bjoerkna* |
| ***Cyprinidae - Complex 4*** | *Alburnus alburnus*<br>*Scardinius erythrophthalmus* |
| ***Lampetra spp.*** | *Lampetra fluviatilis*<br>*Lampetra planeri* |
| ***Leuciscus spp.*** | *Leuciscus idus*<br>*Leuciscus leuciscus* |
| ***Salvelinus spp.*** | *Salvelinus alpinus*<br>*Salvelinus fontinalis* |

### **Appendix S4.** Reference list of the papers retained from the literature research query


Allen, M. C., Kwait, R., Vastano, A., Kisurin, A., Zoccolo, I., Jaffe, B. D., Angle, J. C., Maslo, B., & Lockwood, J. L. (2023). Sampling environmental DNA from trees and soil to detect cryptic arboreal mammals. *Scientific Reports*, *13*(1), 180. https://doi.org/10.1038/s41598-023-27512-8

Bush, A., Monk, W. A., Compson, Z. G., Peters, D. L., Porter, T. M., Shokralla, S., Wright, M. T. G., Hajibabaei, M., & Baird, D. J. (2020). DNA metabarcoding reveals metacommunity dynamics in a threatened boreal wetland wilderness. *Proceedings of the National Academy of Sciences*, *117*(15), 8539–8545. https://doi.org/10.1073/pnas.1918741117

Diana, A., Matechou, E., Griffin, J., Yu, D. W., Luo, M., Tosa, M., Bush, A., & Griffiths, R. A. (2024). eDNAPlus: A Unifying Modeling Framework for DNA-based Biodiversity Monitoring. *Journal of the American Statistical Association*, *120*(549), 1–23. https://doi.org/10.1080/01621459.2024.2412362

Doi, H., Fukaya, K., Oka, S., Sato, K., Kondoh, M., & Miya, M. (2019). Evaluation of detection probabilities at the water-filtering and initial PCR steps in environmental DNA metabarcoding using a multispecies site occupancy model. *Scientific Reports*, *9*(1), Article 1. https://doi.org/10.1038/s41598-019-40233-1

Fukaya, K., Kondo, N. I., Matsuzaki, S. S., & Kadoya, T. (2022). Multispecies site occupancy modelling and study design for spatially replicated environmental DNA metabarcoding. *Methods in Ecology and Evolution*, *13*(1), 183–193. https://doi.org/10.1111/2041-210X.13732

Gold, Z., Sprague, J., Kushner, D. J., Marin, E. Z., & Barber, P. H. (2021). eDNA metabarcoding as a biomonitoring tool for marine protected areas. *PLOS ONE*, *16*(2), e0238557. https://doi.org/10.1371/journal.pone.0238557

Ji, Y., Baker, C. C. M., Popescu, V. D., Wang, J., Wu, C., Wang, Z., Li, Y., Wang, L., Hua, C., Yang, Z., Yang, C., Xu, C. C. Y., Diana, A., Wen, Q., Pierce, N. E., & Yu, D. W. (2022). Measuring protected-area effectiveness using vertebrate distributions from leech iDNA. *Nature Communications*, *13*(1), Article 1. https://doi.org/10.1038/s41467-022-28778-8

McClenaghan, B., Compson, Z. G., & Hajibabaei, M. (2020). Validating metabarcoding-based biodiversity assessments with multi-species occupancy models: A case study using coastal marine eDNA. *PLOS ONE*, *15*(3), e0224119. https://doi.org/10.1371/journal.pone.0224119

McColl-Gausden, E. F., Weeks, A. R., Coleman, R. A., Robinson, K. L., Song, S., Raadik, T. A., & Tingley, R. (2021). Multispecies models reveal that eDNA metabarcoding is more sensitive than backpack electrofishing for conducting fish surveys in freshwater streams. *Molecular Ecology*, *30*(13), 3111–3126. https://doi.org/10.1111/mec.15644

Peixoto, S., Mota-Ferreira, M., Chaves, C., Velo-Antón, G., Beja, P., & Egeter, B. (2023). Multi-species occupancy modeling reveals methodological and environmental effects on eDNA detection of amphibians in temporary ponds. *Environmental DNA*, *5*(4), 796–811. https://doi.org/10.1002/edn3.443

Pukk, L., Kanefsky, J., Heathman, A. L., Weise, E. M., Nathan, L. R., Herbst, S. J., Sard, N. M., Scribner, K. T., & Robinson, J. D. (2021). eDNA metabarcoding in lakes to quantify influences of landscape features and human activity on aquatic invasive species prevalence and fish community diversity. *Diversity and Distributions*, *27*(10), 2016–2031. https://doi.org/10.1111/ddi.13370

Rounds, C. I., Arnold, T. W., Chun, C. L., Dumke, J., Totsch, A., Keppers, A., Edblad, K., García, S. M., Larson, E. R., Nelson, J. K. R., & Hansen, G. J. A. (2024). Aquatic invasive species exhibit contrasting seasonal detectability patterns based on environmental DNA: Implications for monitoring. *Freshwater Biology*, *69*(10), 1479–1493. https://doi.org/10.1111/fwb.14320
